# p75 neurotrophin receptor in pre-adolescent prefrontal PV interneurons promotes cognitive flexibility in adult mice

**DOI:** 10.1101/2023.03.13.532416

**Authors:** Pegah Chehrazi, Karen Ka Yan Lee, Marisol Lavertu-Jolin, Bidisha Chattopadhyaya, Graziella Di Cristo

## Abstract

Parvalbumin (PV)-positive GABAergic cells provide robust perisomatic inhibition to neighboring pyramidal neurons and regulate brain oscillations. Alterations in PV interneuron connectivity and function in the medial prefrontal cortex (PFC) have been consistently reported in psychiatric disorders associated with cognitive rigidity, suggesting that PV cell deficits could be a core cellular phenotype in these disorders. p75 neurotrophin receptor (p75NTR) regulates the time course of PV cell maturation in a cell-autonomous fashion. Whether p75NTR expression during postnatal development affects adult prefrontal PV cell connectivity and cognitive function is unknown. We generated transgenic mice with conditional knockout (cKO) of p75NTR in postnatal PV cells. We analysed PV cell connectivity and recruitment following a tail pinch, by immunolabeling and confocal imaging, in naïve mice or following p75NTR re-expression in pre- or post-adolescent mice using Cre-dependent viral vectors. Cognitive flexibility was evaluated using behavioral tests. PV cell-specific p75NTR deletion increased both PV cell synapse density and the number of PV cells surrounded by perineuronal nets, a marker of mature PV cells, in adult PFC but not visual cortex. Both phenotypes were rescued by viral-mediated re-introduction of p75NTR in pre-adolescent but not in post-adolescent PFC. Prefrontal cortical PV cells failed to upregulated c-Fos following a tail-pinch stimulation in adult cKO mice. Finally, cKO mice showed impaired fear memory extinction learning as well as deficits in a rule set-shifting task. These findings suggest that p75NTR expression in adolescent PV cells contributes to the finetuning of their connectivity and promotes cognitive flexibility in adulthood.

## Introduction

The lack of cognitive flexibility limits an individual’s ability to adapt their goal-oriented behavioral strategies in a changing environment, restricting their choices and making it harder to meet everyday challenges. Numerous studies indicates that the prefrontal cortex (PFC) is critical for cognitive flexibility (1–3). Deficits in cognitive flexibility are the hallmark of prefrontal dysfunction in several psychiatric disorders including schizophrenia, autism spectrum disorder, and intellectual disabilities (4–6). The network and cellular mechanisms underlying cognitive flexibility are currently the focus of intensive studies, especially those using rodent models. A combination of experimental approaches so far suggests that the rodent medial PFC (mPFC) serves as the closest functional homolog to the dorso-lateral PFC in human and non-human primates (7), and is linked to cognitive flexibility in these species.

Within the cortex, there are two main types of neurons: glutamatergic excitatory pyramidal neurons and GABAergic inhibitory interneurons. The most common subtype of GABAergic interneurons, the parvalbumin-expressing interneurons (PV cells), innervate the soma and proximal dendrites of the majority of their neighbouring pyramidal neurons (2, 8–12) and can fire rapid trains of action potentials (13, 14). By virtue of their morphological and electrophysiological properties, PV cells control spike timing in neighboring excitatory cells (15, 16) and modulate the generation of gamma oscillations, which are in turn associated with cognitive flexibility (17–20). In particular, prefrontal cortical PV cell activation facilitates the extinction of cue-based responses (21), a process that can mediate rule-shifts during cognitive flexibility tasks, whereas reduced GABA release from PV cells by targeted *Cacna1a* deletion results in cognitive rigidity (22). On the other hand, however, increasing prefrontal cortical PV cell excitability also impairs cognitive flexibility (5), suggesting that proper PFC circuit function relies on optimally balanced levels of inhibition.

Cortical PV cells undergo extensive synaptic formation, maturation and refinement during the first postnatal month, ultimately acquiring their hallmark exuberant synaptic output towards the end of adolescence (9, 11, 23). Recent studies have started to elucidate the molecular players underlying this unique innervation pattern, either in a cell-autonomous (10, 24–28) or a non cell-autonomous fashion (29–31). In addition to molecular factors promoting PV cell synapse formation, inhibitory mechanisms can also play a role in the fine-tuning of PV cell connectivity (11)Another feature of cortical PV cell maturation is the formation of perineuronal nets (PNN), lattice-like aggregates of chondroitin sulfate proteoglycans-containing extracellular matrix, which embrace their soma and proximal dendrites (32). The aggregation of PNN around PV cells over the first 6 postnatal weeks marks a reduction of PV neuronal plasticity and sets their connectivity (32, 33). We have previously shown that cortical PV cells express the neurotrophin receptor p75 (p75NTR) during the first postnatal month. Conditional knockout of p75NTR in single PV neurons *in vitro* and in PV cell networks *in vivo* caused precocious formation of PV cell perisomatic innervation and perineural nets around PV cell somata in the visual cortex (11, 12), therefore suggesting that p75NTR expression modulates the timing of maturation of PV cell connectivity in the adolescent brain. Whether p75NTR expression in prefrontal cortical PV cells regulates their connectivity in the adult brain, thereby possibly modulating cognitive flexibility is unknown and will be here explored.

## Materials and Methods

## Animals

p75NTR^lox/lox^ mice (Bogenmann et al., 2011), in which exons 4–6 of p75NTR, encoding the transmembrane and all cytoplasmic domains, are flanked by two loxP sites were crossed to the PV_Cre (JAX, B6.129P2-Pvalb ^tm1(cre)Arbr^ /J, 017320, RRID:IMSR_JAX:017320) line to generate PV_Cre^+/-^;p75NTR^lox/lox^ (p75NTR cKO) mice. These lines were maintained through crosses of PV_Cre ^+/-^; p75NTR ^lox/+^ females and p75NTR ^lox/lox^ males to generate PV_Cre^+/-^ ; p75NTR^lox/lox^ and PV_Cre^-/-^ ; p75NTR^lox/lox^ or PV_Cre^-/-^ ; p75NTR^lox/+^ control littermates, or PV_Cre^+/+^; p75NTR^lox/+^ females and p75NTR^lox/+^ males to generate PV_Cre ^+/-^ ; p75NTR^lox/lox^ and PV_Cre ^+/-^ ;p75NTR^+/+^ control littermates, on a mixed 129sv/C57BL/6J background. Cell specificity of Cre-mediated recombination was analyzed by breeding PV_Cre with RCE-EGFP mice (JAX, Gt(ROSA)26Sor^tm1.1(CAG-EGFP)Fsh/Mjax^; Jackson Laboratory, stock #32037), thus generating Ctrl_RCE and cKO_RCE mice. RCE-EGFP mouse line carries a loxP-flanked STOP cassette upstream of the EGFP gene. Removal of the loxP-flanked STOP cassette by Cre-mediated recombination drives EGFP reporter expression. All animals were kept under a light-dark cycle (12 h light–12 h dark) in a temperature and humidity-controlled room. Food and water were available ad libitum. All procedures described here had been approved by the Comité Institutionnel de Bonnes Pratiques Animales en Recherche (CIBPAR) of the Research Center of Sainte-Justine Hospital in accordance with the principles published in the Canadian Council on Animal’s Care’s (Guide to the Care and Use of Experimental Animals.

### Mice genotyping

DNA was extracted from mouse tails and genotyped to detect the presence of Cre alleles and P75NTR conditional and wild-type alleles. Polymerase chain reaction (PCR) was performed using either a set of 2 separated primers (F1 5’-CCTCCGCCAGCTGTCTGCTTCCT -3’ and R1 5’-GGGTGGAAGCTGGGACTGTGCACATGC-3’) to identify the p75NTR floxed versus the wildtype allele or a set of 2 primers (5’-CCTCCGCCAGCTGTCTGCTTCCT-3’, 5’-TCCTCACCCCGTTCTTTCCCC-3’to assure the absence of the p75NTR null allele: with band sizes of 575 bp for the wild-type, 407 bp for the floxed and 400 bp for the null allele. The 3 separate primers used to detect Cre in PV_Cre were: F1(5’-CAGCCTCTGTTCCACATACACTCC-3’), F2(5’-GCTCAGAGCCTCCATTCCCT-3’) and R1(5’-TCACTCGAGAGTACCAAGCAGGCAGGA GATATC-3’) which generate 400 bp and 526 bp (mutant and wild-type) bands. To detect the presence of the RCE allele, 3 separate primers namely, RCE-Rosa1(5’-CCCAAAGTCGCTCTGAGTTGTTATC-3’), RCE-Rosa2 (5’GAAGGAGCGGGAGAAATGGATATG-3) and RCE-Cag3 (5’-CCAGGCGGGCCATTTACCGTAAG-3’) were used, which generated 350bp and 550bp bands.

### Immunohistochemistry

Mice of both sexes were anesthetized, then perfused intracardially with PBS followed by 4% (w/v) paraformaldehyde (PFA) in Phosphate buffer saline (PBS). Intact brains were extracted and post-fixed in 4% PFA/PBS overnight at 4 °C. The tissue was then cryoprotected in 30% (w/v) Sucrose in PBS, sectioned coronally at 40μm on a cryostat (Leica VT100) and stored as floating sections in PBS. For immunohistological analysis, brain sections were blocked in 10% normal goat serum (NGS, Invitrogen, 10000C) in PBS containing 1% (v/v) Triton X-100 for 2 h at room temperature. Primary antibodies were diluted in 5% NGS in PBS containing 0.1% (v/v) Triton X-100 and incubated for 24h-48h at 4 °C. Slices were then washed in PBS (3 x 10’), incubated in the appropriate Alexa-conjugated antibodies in 5% NGS, 0,1% Triton in PBS for 2h at room temperature, washed again in PBS (3 x 10’), and mounted in Vectashield (Vector Lab, H-1000) before imaging. The primary antibodies used in this study and their working concentrations are as follows: mouse monoclonal anti-PV (1:2000; Swant, PV235, RRID:AB_10000343); rabbit polyclonal anti-PV (1:4000; Swant, PV27, RRID:AB_2631173); mouse monoclonal anti-gephyrin (1:500; Synaptic Systems, 147021,RRID:AB_22325461); chicken polyclonal anti-GFP (1:1000; Abcam, 13970, RRID:AB_300798) and rabbit polyclonal anti-cfos (1:500; Synaptic Systems, 226 003, RRID: AB_783431). To label PNNs, a solution of biotin-conjugated lectin Wisteria floribunda (WFA) (10 µg/ml; Sigma-Aldrich, L1516) was added in the primary antibody solution. The secondary antibodies used in this study and their working concentrations are as follows: goat anti-chicken Alexa488 conjugated (1:1000; Abcam, ab150169), goat anti-rabbit Alexa633 conjugated (1:500; Life technologies, A21072), goat anti-mouse Alexa555 conjugated (1:1000; Cell Signaling, 4409S), goat anti-mouse Alexa647 conjugated (1:1000; Cell signaling, 4410S), goat anti-rabbit Alexa555 conjugated (1:400; Life technologies, A21430), and Alexa 568-conjugated streptavidin (1:500; Invitrogen, S-11226). All immunohistological experiments were performed on at least 3 mice per genotype and on 3 different sections per brain region per animal.

### Confocal imaging

All images were acquired using Leica confocal microscopes (SP8 or SP8-STED). We imaged the visual (VCx) and the medial prefrontal cortex (mPFC) using the Leica 20X multi-immersion (NA 0.75) and the 63X oil (NA1.4) objectives. The 20X objective was used to acquire images to analyze the percentage of: GFP^+^PV^+^/PV^+^ cells (recombination rate), GFP^+^PV^+^ /GFP^+^ cells (Cre-recombination specificity), PNN ^+^PV ^+^/PV^+^ cells and PNN^+^PV^+^GFP^+^ /PV^+^GFP^+^ cells. The 63X objective was used to acquire images to analyse perisomatic innervations (number of perisomatic PV^+^/gephyrin ^+^ or Synt2^+^ boutons) and PNN intensity. At least three confocal images from three different brain sections were acquired per brain region with z-step size of 1 µm (20X) or 0.3/0.5 µm (63X). All the confocal parameters were maintained constant throughout the acquisition of an experiment.

### Image analysis

The number and percentages of GFP^+^PV^+^PNN^+^/GFP^+^PV^+^ and PV^+^PNN^+^/PV^+^ was manually quantified using LAS X software (3.3.0.16799) and Neurolucida (MicroBrightField, 11.08.2, 64 bit) For the quantification of PNN fluorescence intensity, all PNN perisomatic rings in each stack were outlined, and the mean gray values were measured after background subtraction, for each experimental condition. Background was determined by measuring mean gray values in at least four different areas (ROI), where immunolabeling was absent, in the same focal plane where PNN perisomatic rings were selected. For the quantification of PNN fluorescence intensity in the AAV treated mice, the PNN mean gray values of the ipsilateral site (ipsi) were normalized to the contralateral site (contra), after subtracting the background as explained above.

To quantify the number of PV^+^ GFP^+^ gephyrin^+^ putative synapses, images were exported as TIFF files and analyzed using Fiji (Image J) software. We first manually outlined the profile of each cell soma and used a series of custom-made macros in Fiji as previously described (Guirado et al., 2018). Briefly, after applying subtract background (rolling value = 10) and Gaussian blur (σ value = 2) filters, the stacks were binarized and the PV ^+^GFP^+^ gephyrin^+^ puncta were independently identified around the perimeter of a targeted cell after selecting the focal plane with the highest soma circumference. At least 4 innervated somata were selected in each confocal image. Puncta were quantified after filtering particles for size (included between 0 and 2 μm^2^) and circularity (included between 0 and 1). All quantification were done by investigators blind to the genotype or experimental conditions.

### Behavioral testing

For all experiments, a camera was mounted above the arena; images were captured and transmitted to a computer running the Smart software (Panlab, Harvard Apparatus) or Freeze Frame software IMAQ 3 (Version 3.0.1.0). The sequence of animals tested was randomized by the genotype. Care was taken to test litters containing both the genotypes specific to the breeding.

### Open Field

Each subject (P60) was gently placed at the center of the open-field arena and recorded by a video camera for 10 min. The recorded video file was later analyzed with the SMART video tracking system (v3.0, Harvard Apparatus). The open field arena was cleaned with 70% ethanol and wiped with paper towels between each trial. The time spent in the center (45% of the surface) versus the periphery was calculated. Locomotor activity was indexed as the total distance travelled (cm).

### Elevated plus maze

The apparatus consisted of two open arms without walls across from each other and perpendicular to two closed arms with walls joining at a central platform. Each subject (P60) was placed at the junction of the two open and closed arms and allowed to explore the maze during 5min while video recorded. The recorded video file was later analyzed with the SMART video tracking system (v3.0, Harvard Apparatus) to evaluate the percentage of time spent in the open arms (open/(open+closed) x100) and the number of entries in the open arms as a measure of anxiety-related behavior.

### Fear conditioning and fear extinction

Fear conditioning and extinction were evaluated in two different contexts (contexts A and B, respectively). The context A consisted of grey walls, a grid floor, and the odor of 70% ethanol during each session. The context B consisted of black and white striped walls, a white plexiglass floor, and the odor of 1% acetic acid during each session. PV_Cre^+/-^; p75NTR^lox/lox^ males and their littermates PV_Cre^-/-^; p75NTR^lox/lox^ or PV_Cre^-/-^; p75NTR^lox/+^ male control mice were conditioned using the following protocol, previously described in (Lavertu-Jolin et al., 2022). Briefly the protocol is as follows, on day 1, mice were allowed to freely explore context A for habituation. On day 2, mice were conditioned in context A by using 5 pairings of the CS (each CS duration of 5secs with white noise at 80 dB), with the co-terminating US (1sec foot-shock at 0.6mA, inter-trial interval: 30–60 s). On days 3 and 4, conditioned mice underwent extinction training in context B where they were presented with 12 pairings of the CS per day (each CS duration of 30secs with white noise at 80 dB). Finally, both retrieval of extinction and context-dependent fear renewal were tested 7 days later in context B and A, respectively, using 4 pairings of the CS (each CS duration of 5secs with white noise at 80 dB).

### Viral vector and stereotaxic injections

pAAV.EF1α-DIO-Flag-p75NTR_T2A_GFP (titre 7.1E12 GC/mL) was cloned from pCIG-p75NTR WT-IRES-eGFP and produced as AAV2/9 serotype by the Canadian Neurophotonics Platform, similar to the pAAV_EF1α_DIO_eYFP control virus (titre 5E13 GC/mL; injected at 1:6 dilution in saline). PV_Cre; p75NTR^+/+^ or PV_Cre; p75NTR^lox/lox^ mice were used at either postnatal day P25 (pre-adolescent age) or >P45 (post-adolescent age) for all surgeries. Unilateral viral injections were performed at the following stereotaxic coordinates measured from the Bregma: +1.9mm Anterior-Posterior, +0.4mm Medio-Lateral and 1.6mm Depth from the cortical surface. Surgical procedures were standardized to minimize the variability of AAV injections. Glass pipette filled with virus was lowered to the right depth at the correct co-ordinates and kept in place for 2 min before starting the injections. A total of 8 injections per mouse at an injection volume of 46nl (total 46×8nl), was performed for each virus. To ensure minimal leakage into surrounding brain areas and to optimise viral spread in the desired area, injection pipettes were kept in the same position following the injection for 5 min. They were withdrawn at the rate of 0.25mm/min following the injection. Mice were kept for 3 weeks following viral injections to allow for optimal gene expression.

### Attentional set shifting task

Behavioral experiments were performed in a home-made acrylic rectangular-shaped maze (30 cm long × 20 cm wide × 18 cm high) divided in half by a guillotine-like door that extended through the width of the maze. One half served as the start area and the other half served as the choice area. In the choice area, an acrylic wall (12 cm long × 18 cm high) presenting four ¼” diameter sniffing holes on the bottom extended out from the back wall and divided the choice area into two equally sized and distinct spatial locations (Supplementary Fig. 1A). Each of these two locations contained a ceramic ramekin identical in color and size (non-porous; depth 3.5 cm × diameter 6 cm). A ramekin filled with water was placed in the starting area.

A piece of Cheerios (approximately 20 mg in weight) was used as the food reward. The cues, either olfactory (odor) or somatosensory and visual (texture of the digging medium which hides the food reward), were altered and counterbalanced (Supplementary Fig. 1B). All odors were essential oils, and unscented digging media. The testing pots were scented a day before to allow the scent to dissipate by adding approximately 0.1 ml of essential oil to the top of the filter paper using a syringe with a 25 G needle.

The experiment was divided into five steps: handling, food restriction, acclimatisation, training, and testing as explained in Heisler et al., 2015. First, mice are single-housed and habituated to a reverse light/dark cycle for at least one week. Mice doing this test were food restricted to 85% of their ad lib. feeding weight in the days prior to testing. When stabilized at 85% of their ad lib. body weight, mice were trained and tested. Mice were on a 85% ad lib food restriction diet during the full duration of the experiment.

### Testing

Mice were tested in seven stages: Simple discrimination (SD), Compound Discrimination (CD), Reversal (Rev), Intra-Dimensional Shift 1 to 3(IDS1, IDS2, IDS3) and Extra-dimensional shift (EDS). Following SD, media and scent were paired for each of the steps. Rewarding ramequin was placed randomly on the left or right side of the maze to avoid the use of spatial cues. Food reward was covered with scented (or not, in the case of SD) media and cereal dust was sprinkled on top of the media to avoid aberrant smell cues. To complete a trial, the mice had to dig into a media to retrieve the food reward. On incorrect trials, animals were free to visit the other side of the apparatus, but the other ramekin was removed. At the end of each trial, animals were guided to the start area and the ramequins were refilled or changed. Animals had 3 min to complete a trial and needed to succeed 8 trials out of 10 to advance to the next stage. The testing phase was scheduled on 2 days. SD, CD, CDR and IDS1 were performed on day 1, followed by IDS2, IDS3 and EDS on day 2.

All mice were handled 2-3 min per day starting 8 days prior to the first day of food restriction and their body weights were recorded each day of handling. Beginning 4 days prior to the start of acclimation, mice were placed on a restricted diet to maintain mice at 80-85% of the *ad libitum* feeding weight by giving 1 gram of food per mouse per day. After mice reached their target weight, they underwent 2 days of acclimation by placing the two ramekins that have been used for food restriction in the testing area of the testing chamber (Supplementary Fig. 1A) with the food reward in each. Food rewards were added continuously to the empty ramekins to encourage the mice to explore the testing area and the pots frequently. The next day after the acclimation (the training day), mice were given 3 min before ending the trial and in subsequent trials, mouse cage bedding was added to the pots. Once the mouse has retrieved the food reward from each pot, it proceeded to the next trial. The trials continue until the food reward is fully covered and the mouse reliably demonstrates the ability to dig in a full pot to find the food reward. Odor and medium combination and location (left or right) of the baited bowl was determined as in (34) (Supplementary Fig. 1B). During each stage of the task, while the particular odor-medium combination present in each of the two bowls may have changed, the particular stimulus that signaled the presence of food reward remained constant over each portion of the task (initial association, rule shift, and rule reversal). In our case, the initial association was paired a specific digging medium with food reward, then the odor was considered the irrelevant dimension. The mouse is considered to have learned the initial association between stimulus and reward if it makes 8 correct choices during 50 consecutive trials. Each portion of the task ended when the mouse met this criterion. If a mouse does not obtain 8 correct choices consecutively within 50 trials, it fails that stage and cannot move on to complete the test. 6 consecutive no choices (3 min without a choice) are considered as a failure to participate and prohibits the mouse from moving on to the next stage.

Following the initial association, the rule-reversal portion of the task began, during which the stimulus that had been consistently not rewarded during the initial association becomes associated with reward. For example, if paper had been associated with reward during the initial association, then during the rule reversal, straw would become associated with reward. Finally, during the extra-dimensional shift stage, the previously negative stimuli is now positive, for example, if felt digging medium was the positive stimuli in the previous stage and paper was the negative stimuli, now the reverse is true. During the extra-dimensional shift stage, the irrelevant dimension (odor in this example) becomes the relevant dimension.

### Tail pinch stimulation protocol

To test whether the effects of p75NTR deletion restricted to PV cells may in turn affect their response following PFC stimulation, a tail-pinch protocol was used. Mice were anesthetized with urethane (1.7 mg/kg, i.p., in sterile saline solution; 94300, Sigma-Aldrich) and 6 tail stimuli were performed, each time by placing forceps pressure on the basal zone of the tail for 15 s, followed by a 3min resting period to recover the basal oscillation pattern. 60 min after the stimulation protocol started, mice were perfused as explained in the histologic procedures section.

### Statistical analysis

All data was analysed using GraphPad Prism 9.3.1. They were first systematically tested for normal distribution with the Shapiro-Wilk normality test. Differences between two groups of normally distributed data with homogenous variances were analyzed using parametric student’s t-test with Welch’s correction, while not normally distributed data were analyzed with the Mann-Whitney test. Fear learning and extinction were analyzed using 2-way Repeated Measure ANOVA, followed by Holm-Sidak *post-hoc* test. To evaluate the effect of the virus-mediated reintroduction on GFP^+^ Gephyrin^+^ perisomatic puncta, we used 1-way ANOVA followed by Tukey’s multiple comparison test. The effect of age and virus treatment on PNN was evaluated by 2-way ANOVA followed by Tukey’s multiple comparison test. The effect of genotype and experimental condition (baseline vs tail pinch) was evaluated by 2-way ANOVA followed by Tukey’s multiple comparison test. Significance level was set to 5% (p = 0.05). Results were considered significant for values of *P*<0.05. Data are presented as mean ± standard error of mean.

### Data availability

All data are available upon request.

## Results

### PV cell-targeted postnatal p75NTR deletion increased PV cell efferent connectivity and PNN aggregation specifically in mPFC of adult mice

Cortical PV cell innervations become progressively more elaborate between the 2^nd^ and 4^th^ postnatal week, with axons forming terminal branches bearing well-defined boutons around the postsynaptic targets (9). To investigate whether p75NTR activity during PV cell postnatal maturation plays a role in PV cell efferent connectivity, we used mice with a conditional allele of p75NTR (*p75NTR^lox/lox^*, (35), which allowed cell-specific developmental stage restricted manipulation of p75NTR, and crossed them to a mouse line with the Cre allele under the control of the PV promoter (PV_Cre, Hippenmeyer PloS Biol PMID 15836427). In cortical GABAergic cells, PV starts to be expressed after the first postnatal week and peaks after the third (12). The above-mentioned cross generated PV-cell restricted homozygous mice (*PV_Cre^+/-^*;*p75NTR^lox/lox^*referred to hereafter as cKO) and their control littermates (*PV_Cre^+/-^;p75NTR^+/+^*referred to hereafter as Ctrl). To unambiguously identify PV cells expressing Cre, we bred these mice with RCE-EGFP mice (JAX *Gt(ROSA)26Sor^tm1.1(CAG-EGFP)Fsh/Mjax^*). At P70-90, we observed that all GFP^+^ cells expressed PV independently of the brain region, and the recombination rate was 98±3% and 69±4% (n=3 mice), in visual cortex (VCx) and mPFC, respectively, which was consistent with previous studies (Amegandjin, Choudhury et al. 2021; Lavertu-Jolin, Chattopadhyaya et al. 2022). We have previously shown that this approach led to significant reduction of p75NTR expression in cortical PV cells (11). To address the role of PV cell-specific p75NTR deletion on their output connectivity, we quantified the putative perisomatic synapses formed by PV cells, identified by the juxtaposition of PV+GFP+ boutons, or GFP+ boutons, and gephyrin, a scaffolding protein present in the postsynaptic sites of GABAergic synapses, both in the mPFC and VCx of P90 cKO mice compared with their control littermates (Fig.1A, B). In our experience, PV immunostaining signal levels could be affected by different factors, such as fixation speed or strength, on the other hand GFP labeling is very stable. For this reason, we quantified both PV+GFP+ boutons or GFP+ boutons. We observed that the density of both perisomatic PV+GFP+gephyrin+ puncta (Fig. 1C) and GFP+gephyrin+ puncta (Fig. 1D) was significantly increased specifically in mPFC, but not VCx, of cKO mice compared to control littermates (Fig. 1E, Fig. 1F). The difference observed in mPFC was not due to changes in the number of putative PV cell presynaptic sites, since the density of perisomatic PV+GFP+ (or GFP+) boutons co-localising with Synaptotagmin 2, which specifically labels PV cell presynaptic sites (36), was not significantly different between the two genotypes (Supplementary Fig.1). Thus, PV cell-specific p75NTR deletion markedly increased the percentage of PV boutons juxtaposed to the postsynaptic marker gephyrin.

**Figure 1.**
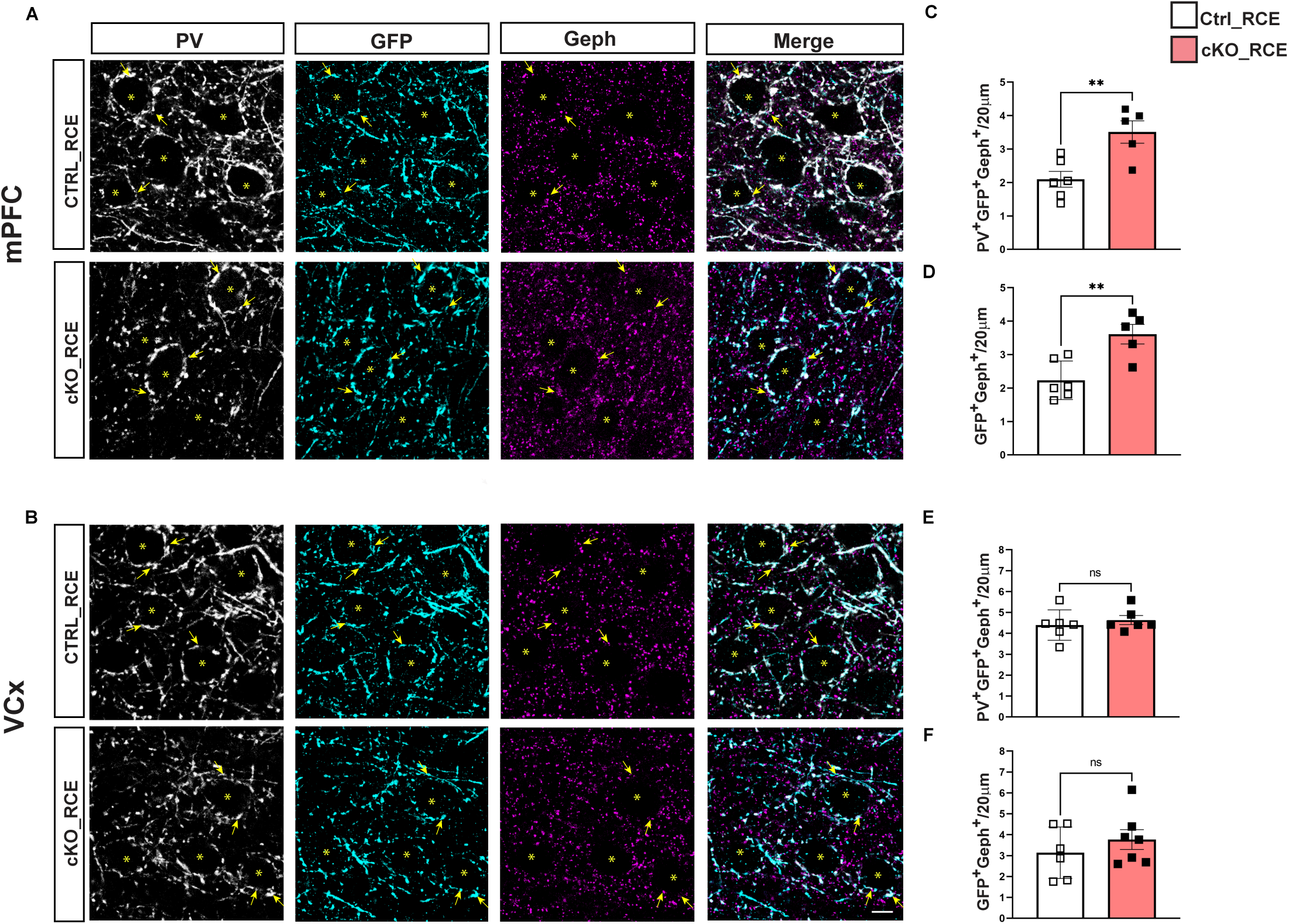
p75NTR deletion in postnatal PV cells leads to increased PV cell innervation in adult mPFC, but not VCx. **A, B**, mPFC (A) and VCx (B) coronal sections of cKO-RCE and Ctrl-RCE littermates labelled with PV (gray), GFP (cyan) and Gephyrin (magenta). Yellow arrows indicate perisomatic PV^+^ GFP^+^ Gephyrin^+^ boutons. Asterisks indicate putative cell somata. Scale bar, 10 µm. **C-F**, PV^+^GFP^+^ Gephyrin^+^ (**C**) and GFP^+^Gephyrin+ (**D**) puncta density is significantly increased (C, Welch’s t test, P=0.009; D, Welch’s t test, P =0.006) in the mPFC, but not in the VCx (**E**, Welch’s t test, P=0.364; **F**, Welch’s t test, P=0.381) of cKO-RCE mice compared to their Ctrl-RCE littermates. cKO-RCE, N=6, Ctrl-RCE, N=5 mice in the mPFC and cKO-RCE, N=6, Ctrl-RCE, N=7 mice in the VCx. Data represent mean and error bars represent ±s.e.m. Open and closed squares represent individual mouse value for *Ctrl*-RCE and cKO-RCE mice, respectively. * P < 0.05, ** P <0.01, *** P <0.001.

PNNs enwrap the somata and proximal dendrites of mature cortical PV cells, stabilizing their afferent synaptic contacts and likely limiting their plasticity (32, 36, 37). Using WFA staining to label PNNs (Fig. 2A,B), we found a significant increase in both PNN immunofluorescence intensity around single PV cell somata and the percentage of PV+GFP+ cells surrounded by PNNs in mPFC of cKO mice compared to Ctrl littermates (Fig. 2C,D). Conversely, only PNN intensity was significantly increased in the VCx of mutant mice (Fig. 2 E, F).

**Figure 2.**
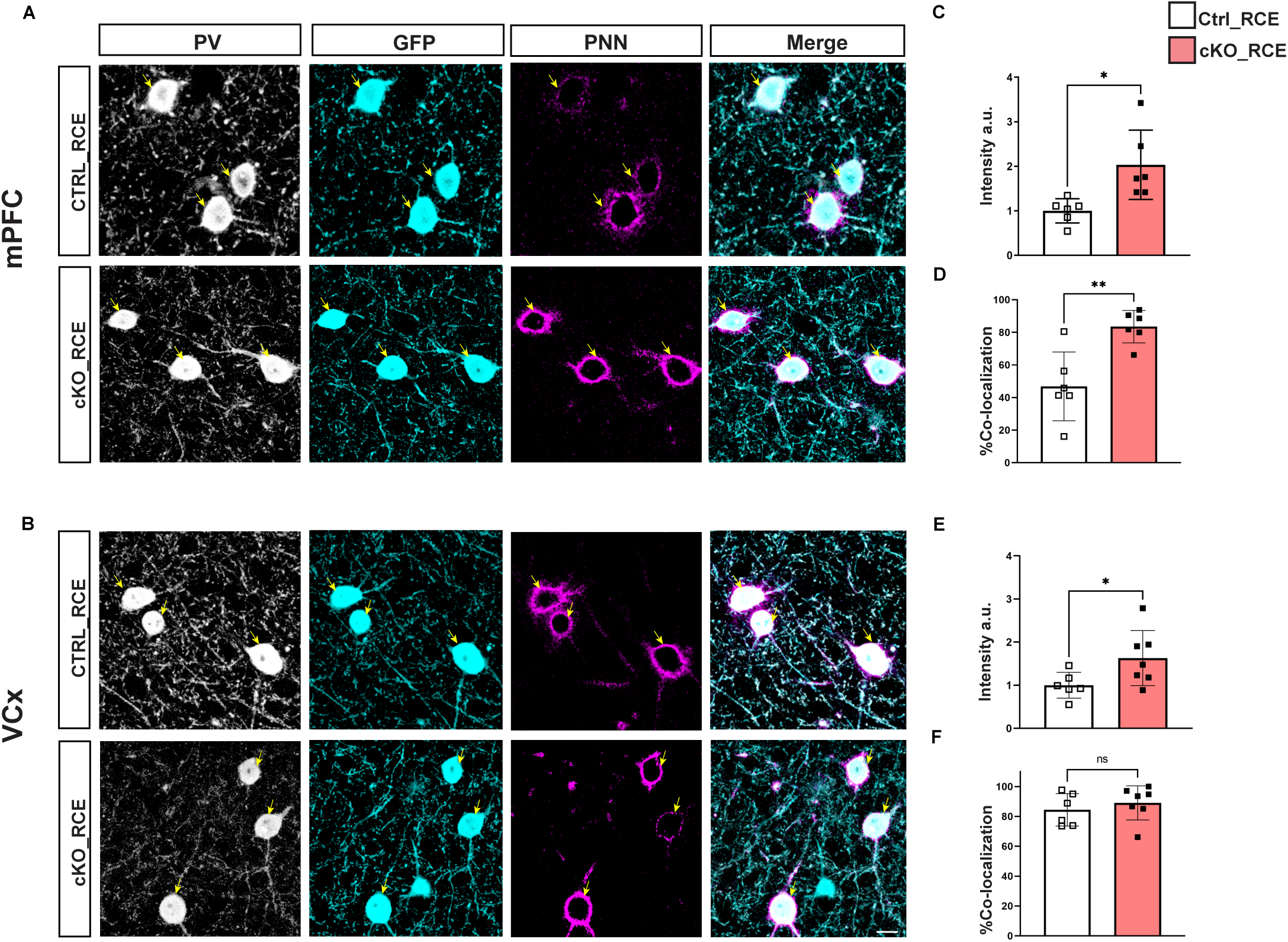
Adult mice with PV cell targeted postnatal p76NTR deletion show increased PNN agglomeration around cortical PV cell somata in mPFC. **A, B,** mPFC (**A**) and VCx (**B**) coronal sections of cKO-RCE and Ctrl-RCE littermates labeled with PV (gray), GFP (cyan) and WFA (PNN, magenta). Yellow arrows indicate PV^+^GFP^+^PNN^+^ cell bodies. Scale bar, 10 µm. **C-F**, Mean PNN intensity (**C**) and the proportion of PV^+^ cell somata surrounded by PNN (**D**) are significantly increased in the mPFC of cKO-RCE mice compared to their littermates (**C**, Welch’s t-test, P=0.021; D, Welch’s t-test, P=0.006). In the VCx, only PNN intensity is significantly increased (**E**, Welch’s t-test, P =0.045; F, Welch’s t-test, P=0.469). Ctrl-RCE, N=6 for both regions, cKO-RCE, N=7 mice in VCx and N=6 mice in mPFC. Data represent mean and error bars represent ±s.e.m. Open and closed squares represent individual mouse value for Ctrl-RCE and cKO-RCE mice, respectively. * P < 0.05, ** P <0.01, *** P <0.001.

Our data therefore suggests that p75NTR, in postnatal prefrontal cortical PV cells, constrains both their synaptic innervation and the aggregation of PNNs around their somata.

### AAV mediated reintroduction of p75NTR in pre-adolescent, but not post-adolescent, prefrontal PV cells rescues their synaptic output and PNN intensity

PV cell hyper-connectivity and increased PNN expression around their somata in adult conditional p75NTR knockout mice could be directly caused by p75NTR signaling dysregulation in PV cells or induced as a consequence of brain-wide homeostatic feedback mechanisms that influence neural circuit development. To distinguish between these two possibilities, we virally re-expressed p75NTR in the mPFC of *PV_Cre^+/-^;p75NTR^lox/lox^* and *PV_Cre^+/-^;p75NTR^+/+^*mice using an inverted double-floxed p75NTR coding region driven by a ubiquitous EF1a promoter (Fig. 3A). To determine whether p75NTR expression in PV cells affected the refinement of their connectivity during adolescence (11, 23), or whether it played a role in the maintenance of their innervation fields after they reached maturity, we conditionally reintroduced it in PV cells either at P25 (pre-adolescence) or >P45 (post-adolescence) (Fig. 3B). Since PV expression levels could be affected by AAV injection, we ascertained that the density of perisomatic boutons identified using Cre-dependent GFP expression was similar to the density quantified using both PV and GFP (Fig.1C-G, and Supplementary Fig.1), and therefore relied on GFP expression to identify putative PV cell boutons. Consistent with our previous data (Fig. 1D), we observed a significant increase in the density of putative PV cell synapses, identified by the co-localisation of GFP and gephyrin around pyramidal neurons, in cKO mice compared to control mice when both were injected with the control virus (AAV_DIO_YFP) during either pre-adolescence or post-adolescence (Fig. 3D). Injection of AAV_DIO_p75NTR-T2A-YFP mice at P25, allowing Cre-dependent p75NTR re-expression, was sufficient to reduce the density of perisomatic GFP+ Gephyrin+ puncta to Control levels in adult cKO. The same treatment in post-adolescent mice however had no such significant effects (Fig. 3D). We then investigated whether p75NTR reintroduction could also affect PNN aggregation around PV cells, by comparing the intensity of PNN staining around PV cell somata in the injected (ipsilateral) versus the non-injected (contralateral) mPFC within the same mouse. In this case, a ratio of 1 would indicate no difference in PNN intensity in the ipsi-versus the contralateral mPFC, while a ratio <1 would indicate reduced PNN intensity in the injected, ipsilateral site. In *PV_Cre^+/-^ ;p75NTR^+/+^* mice transduced with AAV_DIO_YFP (control mice) we found a ration of ∼ 1 (1.1 ±0.2, n=4 mice), suggesting that viral injection per se did not affect PNN intensity. Pre- or post-adolescence cKO mice were then injected with either AAV_DIO_YFP or AAV_DIO_p75NTR-T2A-YFP. We found that transduction of p75NTR in PV cells was sufficient to decrease the intensity of PNN enwrapping them, only when re-introduced in pre-adolescent, but not in post-adolescent mice (Fig. 4).

**Figure 3.**
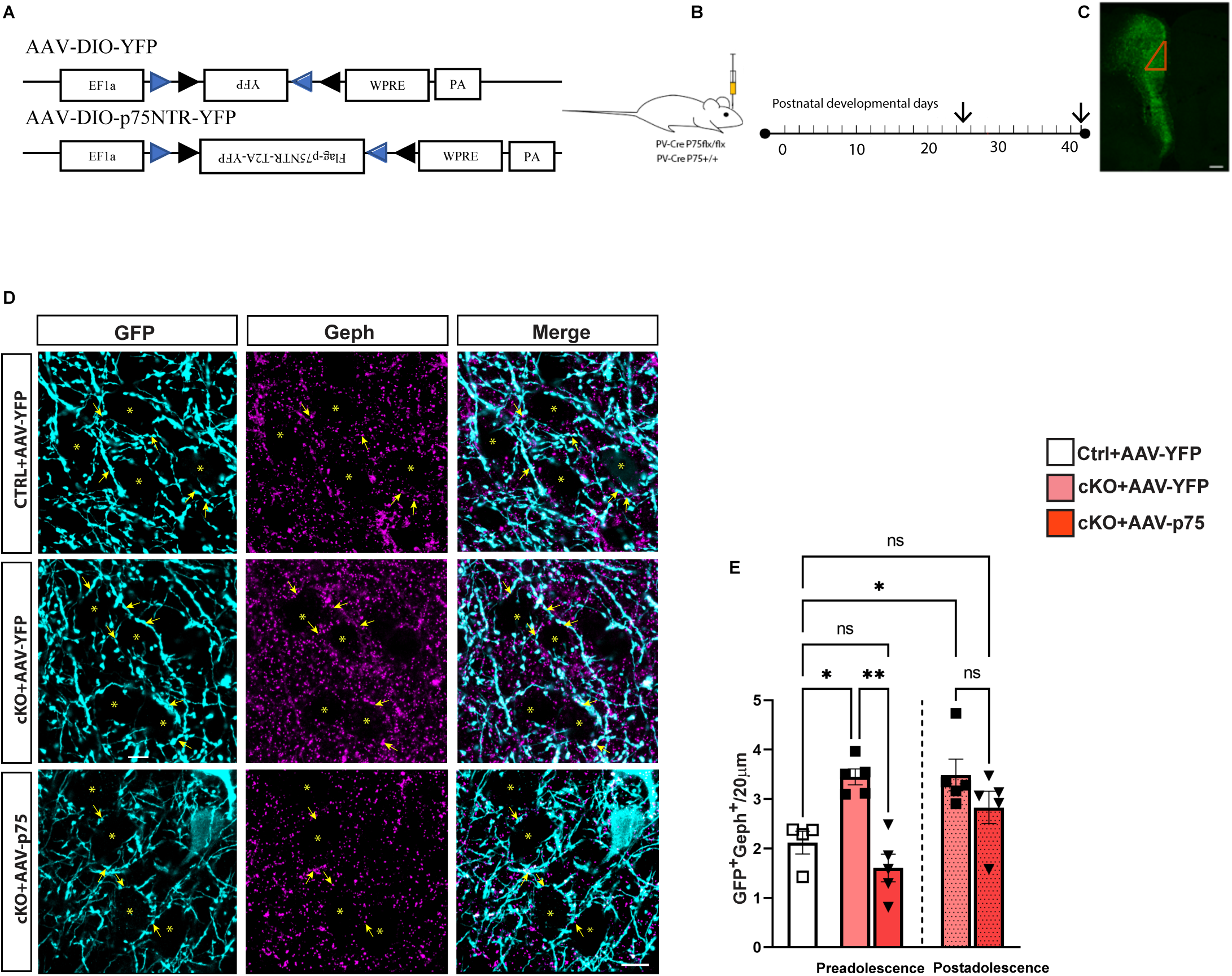
PV-cell specific p75NTR re-expression in pre-adolescent, but not post-adolescent, cKO mice normalizes mPFC PV cell efferent connectivity. **A**, Schematic representation of Cre-dependent viral vectors. **B**, Time line of viral vector injections. **C**, Coronal section of mPFC representative of viral vector transduction shown by GFP immunolabeling (green). The red triangle indicates the prelimbic cortex. **D**, Immunolabelling of GFP (cyan) and Gephyrin (magenta) in mPFC coronal sections from *PV_Cre^+/-^; p75NTR^+/+^* (Ctrl) and *PV_Cre^+/-^; p75NTR^lox/lox^*(cKO) mice injected with Cre-dependent YFP expressing virus (AAV_DIO_YFP, indicated as AAV-YFP) and from *PV_Cre^+/-^; p75NTR^lox/lox^* injected with the Cre-dependent p75NTR expressing virus (AAV_DIO_p75NTR_T2A_YFP, indicated as AAV-p75). Yellow arrows point to GFP+ Gephyrin+ colocalized puncta. Asterisks indicate putative cell somata. Scale bar, 10 µm. **E**, Mean perisomatic GFP^+^Gephyrin^+^ bouton density is significantly increased in cKO + AAV-YFP compared to Ctrl+AAV-YFP mice (one-way Anova with Tukey’s multiple comparison test, F=9.156; P=0.0003; Ctrl+AAV-YFP vs pre-adolescent injected cKO mice: P= 0.028; Ctrl+AAV-YFP vs post-adolescent injected cKO mice, P= 0.029). GFP^+^Gephyrin^+^ puncta density is decreased in cKO + AAV-P75 compared to cKO + AAV-YFP mice injected at P25 (P= 0.001), while it is not significantly different from the values of CTRL +AAV-YFP mice (ns, P= 0.712). cKO + AAV-p75 and cKO+ AAV-YFP mice injected after the end of adolescence do not show significant difference in GFP^+^ Gephyrin^+^ puncta density (P= 0.445). Number of mice: Ctrl+AAV-YFP, n=4, cKO + AAV-YFP, n=5 injected as preadolescent and n=5 injected as post-adolescent, cKO+ AAV-p75, n= 5 injected at P25 as preadolescent and n=5 injected as post-adolescent. Data represent mean ± s.e.m. Symbols represent individual mouse value. * P < 0.05, ** P <0.01, *** P <0.001. ns= non significative.

**Figure 4.**
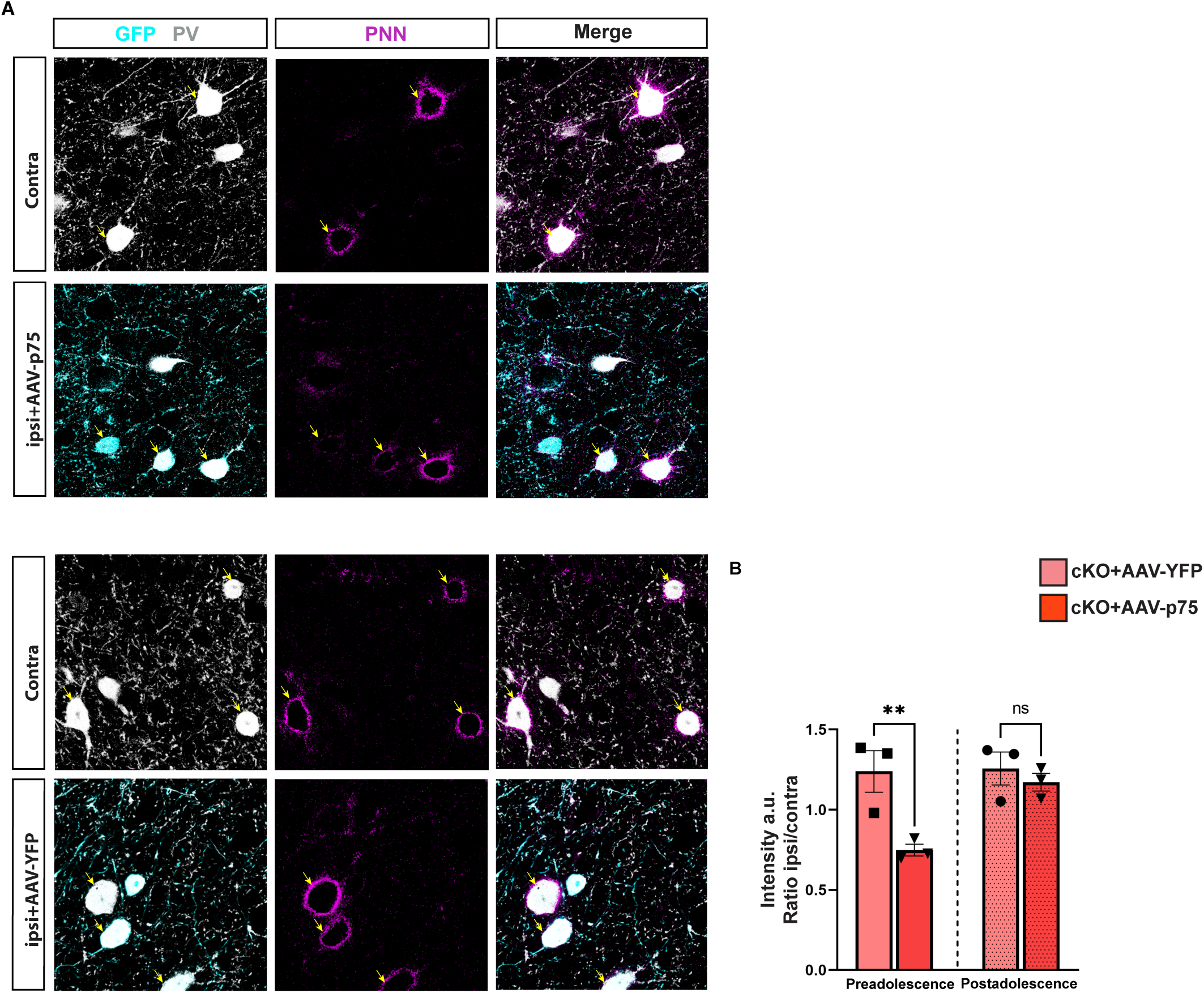
PV-cell specific p75NTR re-expression in pre-adolescent, but not post-adolescent, cKO mice reduces PNN intensity. **A**, GFP (cyan) and WFA (PNN, magenta) labeling in the three experimental groups. Yellow arrows indicate GFP^+^ cell somata surrounded by PNN. Scale bar, 10 µm. **B**, Ipsi/contra PNN intensity around PV cell somata is significantly lower in cKO mice injected with AAV p75 compared to cKO injected with AAV-YFP at P25 (two-way Anova; F_treatment_ (1,8)=10.45, P=0.012; F_age_ (1,8) =6.122, P=0.038, F_treatment*age_ (1,8)=5.145, P=0.053 Sidak’s multiple comparison test; cKO+AAV-YFP vs cKO+AAV-p75, P= 0.009), but not in mice injected after the end of adolescence (cKO+AAV-YFP vs cKO+AAV-p75, P= 0.764). Number of mice: Ctrl+AAV-YFP n=4, cKO+AAV-YFP n=3 injected at P25 and n=3 injected after adolescence, cKO+AAV-p75, n= 3 injected at P26 and n=3 injected after adolescence. Data represent mean ± s.e.m. Symbols represent individual data points. * P < 0.05, ** P <0.01, *** P <0.001. ns= non significative.

Altogether, these data strongly suggest that p75NTR in mPFC PV cells contribute to the fine-tuning of their connectivity and to the aggregation of PNN around their somata during adolescence.

### PV cell-specific p75NTR removal affects their engagement following a tail pinch stimulation protocol

So far, we observed that mice carrying PV cell-specific postnatal p75NTR deletion showed significant increase in PNN aggregation specifically around mPFC PV cells (Fig.2). Several lines of evidence suggest that the PNN around PV cells affects their cellular and synaptic plasticity, thus modulating their recruitment in local networks (32, 33, 36, 38, 39). To test whether the observed increase in PNN affects PV cell recruitment in the *PV_Cre^+/-^;p75NTR^lox/lox^* mice, we next analyzed the expression of the cellular activity marker c-Fos in PV cells following a tail-pinch protocol, which is known to induce a synchronous activated state in prefrontal neuronal circuits (39, 40). During this protocol, the tail was stimulated 6 times by placing pressure on the basal zone with forceps for 15 sec, followed by a 3 min resting period (Fig. 5A). Mice were then perfused 45 min after the last tail pinch. We observed a significant increase in the intensity of c-Fos expression in PV cells from control mice following the tail pinch stimuli compared to PV cells from non-pinched mice. However, no significant increase in cFos was observed in cKO mice (Fig. 5 B, C), suggesting decreased PV neuronal network engagement in the absence of p75NTR.

**Figure 5.**
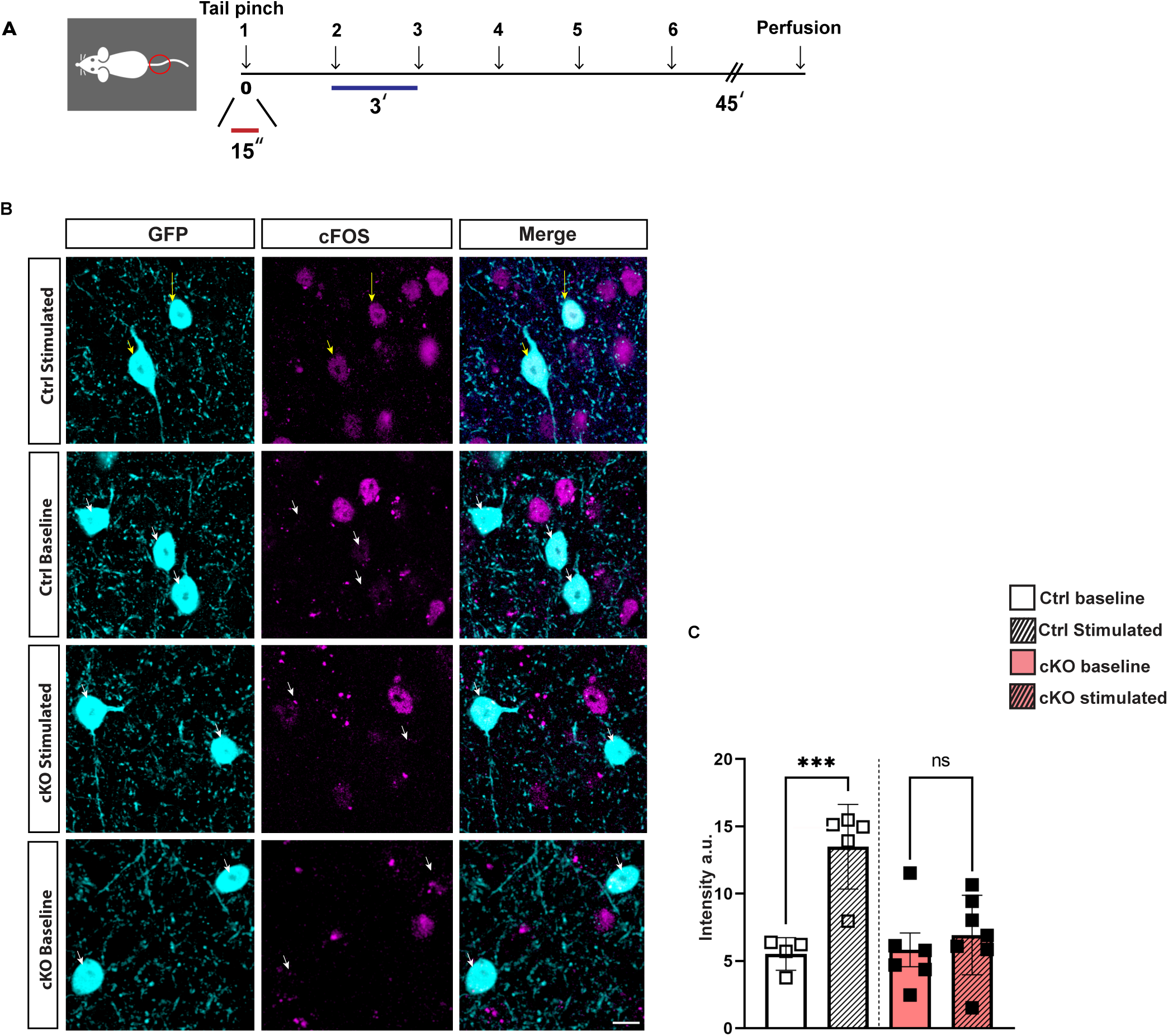
Prefrontal cortical PV cells from cKO mice do not show c-Fos increased expression following a tail pinch stimulation. **A**, Schematic representation of tail pinch stimulation experimental approach. **B**, Immunolabelling of GFP (cyan) and c-Fos (magenta) in mPFC coronal sections of cKO-RCE and Ctrl-RCE littermates without or following sensory stimulation. Yellow arrows indicate c-fos positive nuclei in GFP+ cells, while white arrows indicate c-Fos negative nuclei in GFP+ cells. Scale bar, 10 µm. **C**, Graph showing the fluorescence intensity of c-Fos in PV neurons. Ctrl mice showed significantly higher c-Fos expression in GFP+ cells following tail pinch stimulation than at baseline condition (two-way ANOVA; F_genotype_ (1,18) =6.433, P=0.02, F _experimental condition_ (1,18)=13.55, P=0.001, F_experimental condition*genotype_ (1,18)=7.792 P=0.012 Tukey’s multiple comparison test, P=0.001), while cKO mice do not show any significant difference (ns, P=0.756). Data represent mean ± s.e.m. Ctrl-RCE: Baseline n=4, stimulated n= 5; cKO-RCE baseline n=6, stimulated n=7. * P < 0.05, ** P <0.01, *** P <0.001. ns= non significative.

### Adult *PV-Cre; p75^lox/lox^* mice exhibit impaired fear extinction learning and cognitive flexibility

Altogether, our data shows a region-specific effect of PV cell-restricted postnatal p75NTR deletion in prefrontal cortical PV cells compared to visual cortical PV cells. In mPFC, PV cell activity modulates behaviors linked to cognitive flexibility, such as the extinction of a learnt cue-reward association (21),or learning new associations in a set-shifting task (5, 34). Therefore, we next sought to understand whether cognitive flexibility was affected in the cKO mice, by analysing extinction of cue-mediated aversive behavior, a form of cognitive flexibility dependent on the mPFC (41, 42), as well as behavioral performance in a set-shifting-task.

Since extinction of a learned behaviour is a form of cognitive flexibility, we first used fear extinction as a behavioral assessment tool and analysed fear memory acquisition, extinction rate and spontaneous recovery in cKO mice as compared to control littermates (Fig. 6A). During fear conditioning, both cKO and control mice presented strong freezing behaviour, suggesting that fear learning was not affected by p75NTR deletion in PV cells (Fig. 6B). On the first day of extinction training (early extinction, CS 1-6; Fig.6C), cKO mice showed freezing times similar to control littermates. However, the difference became apparent on the second day of extinction training (late extinction, CS 12; Fig.6C) where the cKO stayed at higher freezing times as compared to control littermates. The difference in freezing persisted during the retrieval test 7 days later (Fig.6D), suggesting a lack of extinction learning and stronger recovery of fear memory. During the renewal test, both control and cKO mice showed similar freezing (Fig.6E), suggesting that extinction memory was not generalized to different contexts in control mice, and that cKO mice did not generate stronger memories compared to their control littermates.

**Figure 6.**
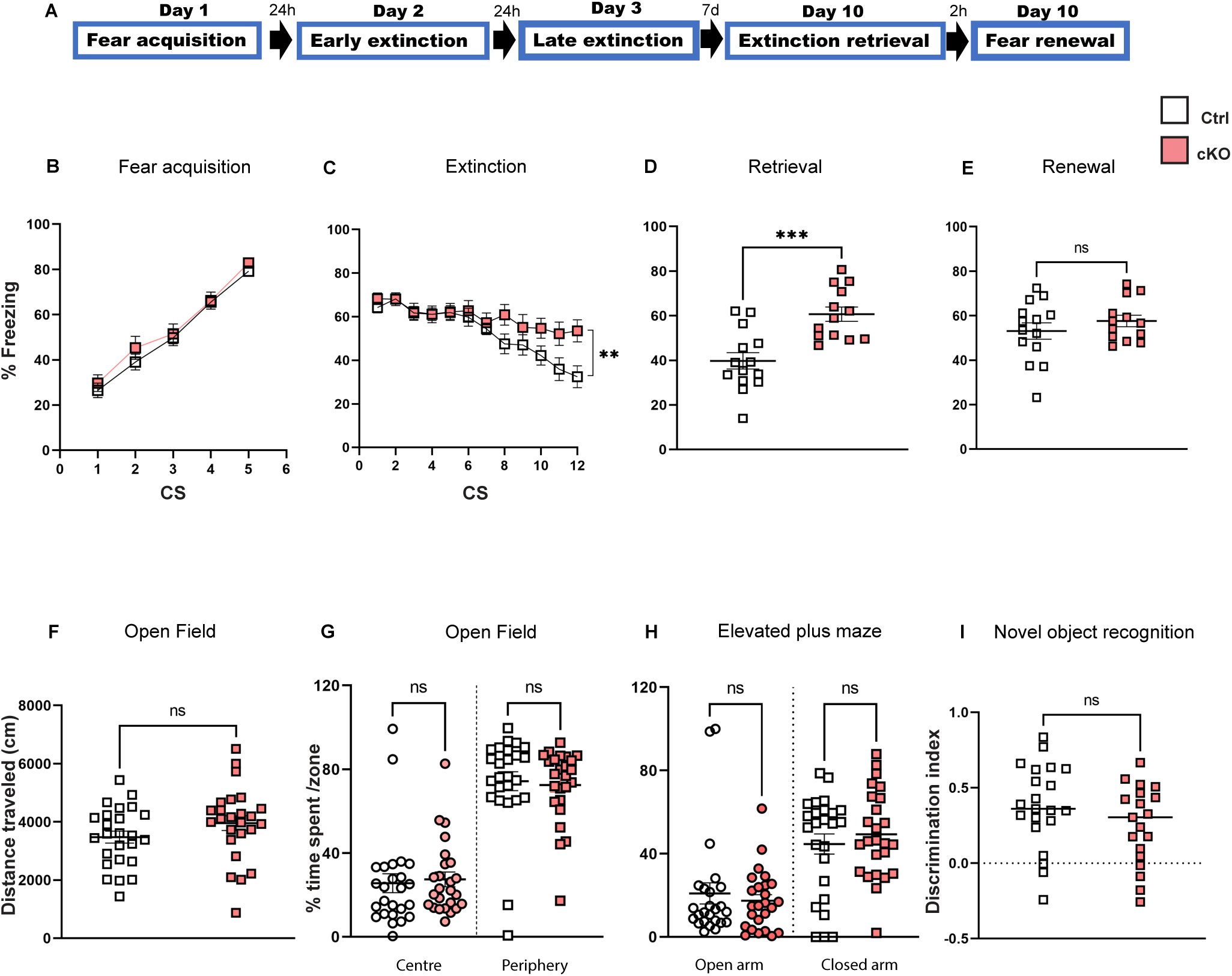
Adult mice with PV cell-restricted postnatal p75NTR deletion show impaired fear extinction learning, but normal locomotion, anxiety behavior and novel object recognition memory. **A**, Schematic representation of the experimental protocol for fear memory acquisition, extinction, and spontaneous recovery. **B-D**, cKO mice show no difference in fear acquisition compared to their control littermates (**B**, two-way Repeated Measure ANOVA; F_genotype_ (1,25) =0.6815, P=0.4169, F_CS_ (4,100) =148.2, P< 0.0001, F_genotype*CS_ (4, 100), P=0.8044), but impaired fear extinction learning (C, two-way Repeated Measure ANOVA; F_genotype_ (1,25)=2.402, P=0.133, F_CS_ ( 11, 275), P< 0.0001, F_genotype*CS_ (11, 275), P=0.0003, Sidak’s post-hoc test; P=0.0043 at CS12) and increased spontaneous recovery of fear memory at the fear retrieval test (**D**, Welch’s t test, P=0.002). **E**, The two genotypes show no difference in fear renewal test (Welch’s t test P=0.322). Number of mice: *P75NTR^lox/lox^*n=14; *PV-Cre; p75NTR^lox/lox^* n=13. **F-H**, cKO and control mice travelled similar distance (**F**, Welch’s t test, open field, P= 0.147) and spend similar time in the center and in the periphery (**G**, multiple unpaired t-test, center: P= 0.748, periphery: P=0.748) of a novel open field and the closed and open arms of the elevated plus maze (**H**, multiple unpaired t-test closed arms: P=0.471, open arms: P= 548). n = 25 mice per genotype. **I**, cKO and control littermates show no difference in the novel object recognition (NOR) task (discrimination index, Welch’s t test, P=0.147). Ctrl, N = 21, cKO, N=19 mice. Data represent mean ± s.e.m. Number of mice: *P75NTR^lox/lox^* n=21; *PV-Cre; p75NTR^lox/lox^* n=19. CS= conditioned stimulus. * P < 0.05, ** P <0.01, *** P <0.001. ns= non significative.

The lack of extinction learning was not due to altered locomotion or increased anxiety since cKO mice did not show significant differences in distance covered (Fig.6F) and the time spent in the center versus the periphery in the open field (Fig.6G) or the open versus closed arms in the elevated plus maze (Fig.6H). Further, cKO mice did not show alterations in memory function measured by the novel object recognition (NOR) task (Fig.6I). Overall, these data suggest that PV cell-restricted postnatal p75NTR deletion leads to specific deficits in fear extinction learning, which is a form of cognitive flexibility.

To further support this finding, we used the attentional set shifting task (Fig.7A), which is equivalent to the Wisconsin Card Sorting Task (WCST) used in humans to assess prefrontal cortex mediated cognitive flexibility (3). During this test, the mice were presented with two bowls on each trial and must choose to dig in one bowl to find the food reward. Each bowl contained a different odor and a different digging medium, and the odor-medium combinations varied from trial to trial. During the initial association phase, the food restricted mouse learnt to associate the relevant cue (*i.e.,* a specific digging medium) and ignore an irrelevant cue (*i.e.,* odor), by pairing a food reward with the medium (initial association, simple discrimination SD, followed by compound discrimination CD; Fig. 7A). During the reversal stage, the mouse had to reverse rules and learn that the food was now associated with the digging medium that had not been rewarded during the initial association (Rev1, Fig. 7A). The stimulus-reward association was then reinforced in subsequent tasks where the type of digging medium and odor changed, but the paired association between medium and reward remained (intra-dimensional shift, IDS1, IDS2, IDS3, Fig. 7A). During the extra-dimensional shift stage, the irrelevant dimension (odor for example) became the relevant dimension (EDS, Fig. 7 A). We observed that while both control and cKO mice were able to learn the initial association in a similar number of trials, an increasing number of cKO mice failed to modify their response when the rules changed during different stages of the test (Fig.7B). Taken together, the behavioral data supports the conclusion that PV cell-specific deletion of p7NTR in pre-adolescent mice leads to cognitive flexibility deficits in adulthood.

**Figure 7.**
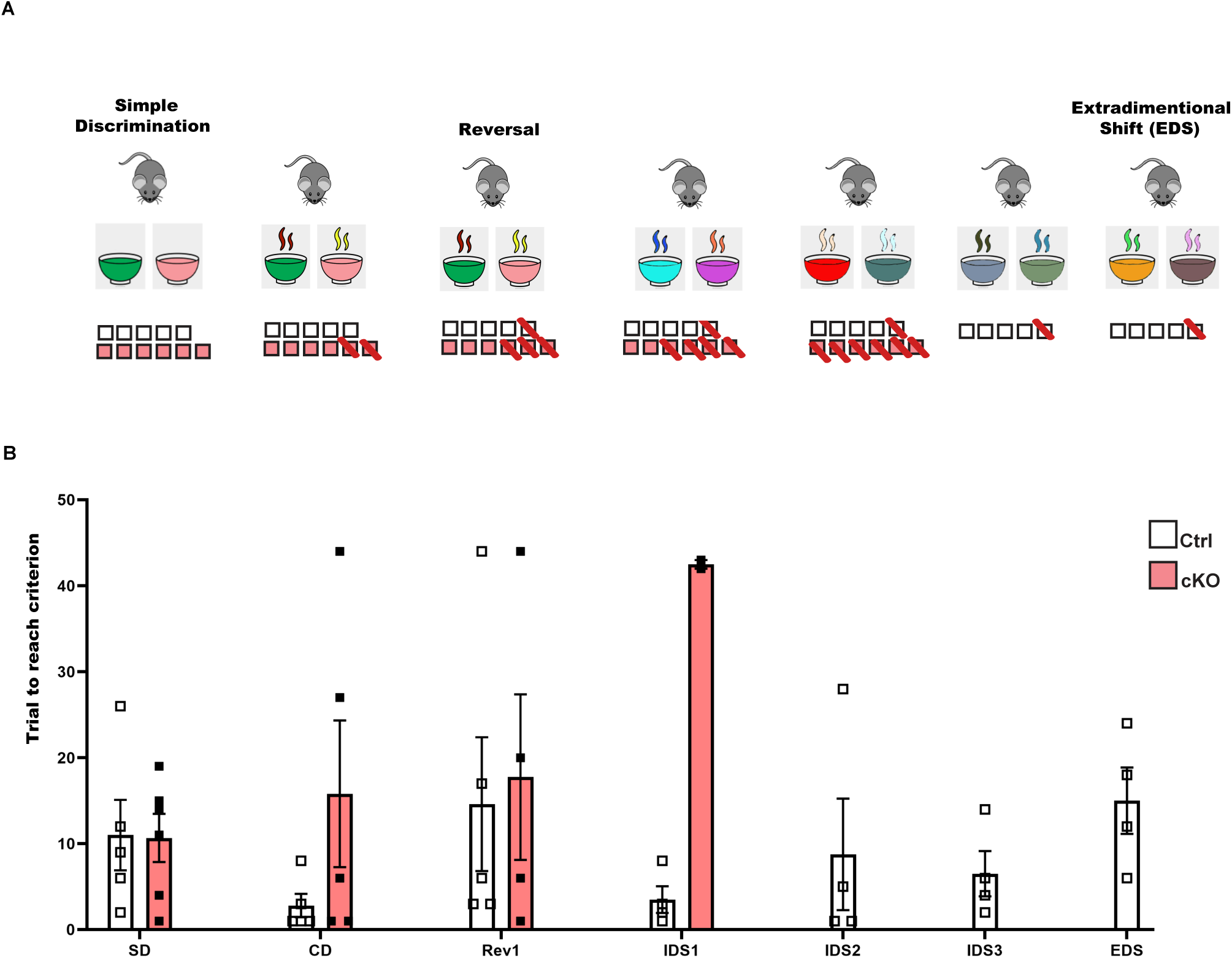
Adult mice with PV cell-restricted postnatal p75NTR deletion show impaired cognitive flexibility. **A**, Attentional set-shifting task protocol. **B**, Average number of trials to meet criterion (eight consecutive correct trials) for each discrimination. The number of mice that succeeded at each stage of the test are represented with an open symbol, while those that failed are crossed out with a red line. Data represent mean ± s.e.m. Ctrl, N = 5, cKO, N=6 mice.

## Discussion

In this study, we investigated whether and how p75NTR expression in PV cells during the juvenile and adolescent period regulates their mature phenotype in adulthood. We found that PV cell-restricted p75NTR deletion leads to increased PV cell connectivity and PNN aggregation in the prefrontal, but not visual cortex in adulthood. We further showed that these mice showed impaired fear memory extinction learning as well as deficits in a complex rule set-shifting task, which are behavioral alterations suggestive of cognitive rigidity.

Altogether, these data suggest that p75NTR expression in mPFC PV cells acts as a negative signal constraining the formation of PV cell connectivity in adult mice. This effect is restricted to the juvenile and adolescent period since prefrontal cortical PV cell hyperconnectivity could be reversed by viral-mediated p75NTR reintroduction in pre-but not post-adolescent mice. p75NTR activation may constrain the formation of PV cell innervation locally by regulating cytoskeletal dynamics (43–45) or sensitizing neurons to other inhibitory, growth cone collapsing cues (46–48). p75NTR-mediated signaling may also cause changes in the synthesis of specific proteins, including those required for PNN condensation around PV-positive cells (33).

Consistent with a specific role of p75NTR expression in juvenile and adolescent prefrontal cortex, we did not previously find significant differences in either PV cell perisomatic connectivity or visual functional properties in visual cortex of adult *PV_Cre;p75NTR^lox/lox^* mice ((11). One possible explanation is that the removal of p75NTR protein in *PV_Cre;p75NTR^lox/lox^* mice occurs too late to affect the maturation of visual cortical circuits, which have for the most part plateaued by the first postnatal month (49). Conversely, PFC circuitry continues to mature through adolescence into young adulthood. Indeed, adolescence is a particularly sensitive time for PFC maturation as environmental stimuli during this period can elicit long-lasting effects on PFC function and behavior (50–53).

Another, non-mutually exclusive possibility is that the availability of p75NTR ligand(s) may be different in distinct cortical regions during adolescence, leading to region-specific recruitment of p75NTR signaling. The role of neurotrophins, their precursor forms (pro-neurotrophins) and prodomains in p75NTR-mediated signaling has been the subject of a multitude of studies (54–57). While it is well accepted that the prodomain plays a role in the folding, stability, and intracellular trafficking of BDNF (58), recent data suggest that the prodomain per se may have diverse biological functions (57, 59, 60). During the transition from childhood to adolescence, BDNF levels increase significantly in the frontolimbic circuitry (61). Cleaved BDNF prodomain can also be secreted in an activity-dependent manner from neurons and peaks at peri-adolescence in the hippocampus (57). Of note, human and mouse carriers of a common single nucleotide polymorphism in the *BDNF* gene resulting in a Val66Met substitution in the prodomain region have impaired fear extinction learning, decreased responses to fear extinction-based therapies and enhanced risk of developing depression and anxiety disorders (62–64). Whether BDNF prodomain levels and/or neuron subtype-specific localisation of p75NTR and related signaling molecules are different in prefrontal versus sensory cortical areas in juvenile and adolescent mice remains to be explored.

p75NTR deletion in postnatal PV cells also led to a significantly higher number of PV cells enwrapped by PNN in adult mPFC. A recent study found that in adult mice, prefrontal cortical PV cells enwrapped by PNN show higher density of perisomatic glutamatergic and GABAergic boutons when compared with PV cells lacking PNN (39). Further, PNN enzymatic removal has been shown to reduce γ oscillations, which are particularly dependent on PV cell activity (65, 66), in the prelimbic region of PFC following a tail-pinch in anesthetized mice (39). A similar treatment led to reduced inhibitory neuron spiking activity recorded from the visual cortex of freely moving adult rats (67). The specific effects of increased PNN density on PV cell synaptic drive remain to be explored; nevertheless, we observed that adult PV cells in the mPFC of *PV_Cre;p75NTR^lox/lox^* mice showed no increase in cFos expression following a tail-pinch sensory stimulation, suggesting that their recruitment in local circuits might be impaired. Of note, both reducing GABA release from PV cells (34) or increasing prefrontal cortical PV cell excitability impairs cognitive flexibility (5), suggesting that proper PFC circuit function relies on balanced levels of inhibition provided by PV cells.

All together these findings suggest that alterations of molecular factors regulating the timing of PV cell maturation in the juvenile brain can lead to lasting consequences on the connectivity these cells make within cortical circuitry, thus contributing to long-term cognitive impairments. Furthermore, PV cells in slow maturing circuits, such as those in mPFC, are likely to be more sensitive to environmental or pharmacological-dependent induced alterations in molecular signaling, thus creating a window of vulnerability that could contribute to the development of cognitive symptoms in adulthood.

## Acknowledgements

We would like to thank Antônia Samia Fernandes do Nascimento for technical assistance, Vidya Jadhav, Maude Larouche and Menna Khaled for help with data analysis. We thank the Canadian Neurophotonics Platform for producing the viruses used in our studies. We acknowledge the Comité Institutionnel de Bonne Pratiques Animales en Recherche (CIBPAR), and all the personnel of the animal facility of the Research Center of CHU Sainte-Justine (Université de Montreal), as well as Dr. Elke Küster-Schöck at the plateforme d’imagerie microscopique (PIM) for instrumental technical support.

## Author contributions

PC, BC and GDC designed the experiments. PC, KL, BC performed the experiments. PC, KL and BC analysed data. M L-J provided critical reagents and technical support. PC, BC and GDC wrote the manuscript. All authors read and corrected the manuscript.

## Legends

**Supplementary Figure 1. Experimental setup for attentional set shift-task. A**, Maze. **B**, Different digging medium used during the test.

**Supplementary Figure 2. The number of putative PV cell presynaptic sites is not significantly affected by postnatal p75NTR deletion in PV cells.** A, B. mPFC (A) and VCx (B) coronal sections of cKO-RCE and Ctrl-RCE littermates labelled with GFP (cyan), Synaptotagmin 2 (Syn2, magenta) and PV (gray). Yellow arrows indicate perisomatic PV+ GFP+Syn2 + boutons. Asterisks indicate putative cell somata. Scale bar, 10 µm. C-F, PV + GFP+ Syn2+ (C-E) and GFP+ Syn+ (D-F) puncta densities are not significantly different between the two genotypes neither in the mPFC (C, Welch’s t test, P=0.696; D, Welch’s t test, P=0.965) nor in the VCx (E, Welch’s t test, P=0.493; F, Welch’s t test, P value=0.445). cKO-RCE, N=4, Ctrl-RCE, N=5 mice. Data represent mean and error bars represent ±s.e.m. Open and closed squares represent individual mouse value for *Ctrl*-RCE and cKO-RCE mice, respectively. ns= non significative.

