## Supplementary figures and images for "p75 neurotrophin receptor in pre-adolescent prefrontal PV interneurons promotes cognitive flexibility in adult mice"

### Supplemental Figures

Supplementary figure 1

A

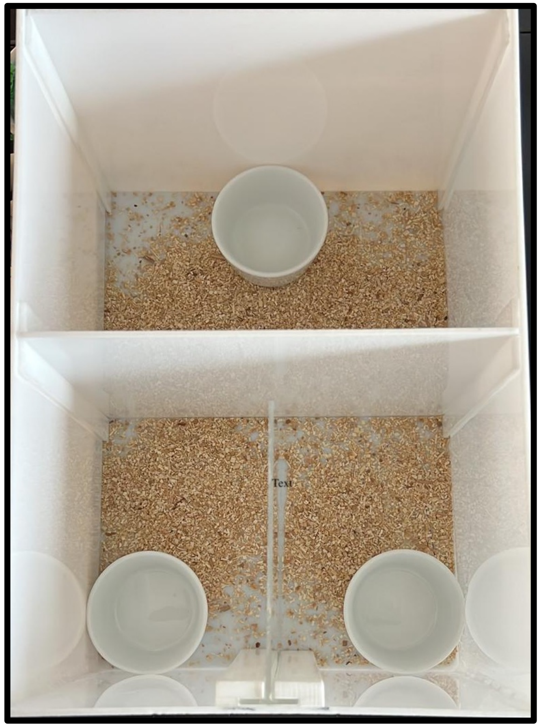

B

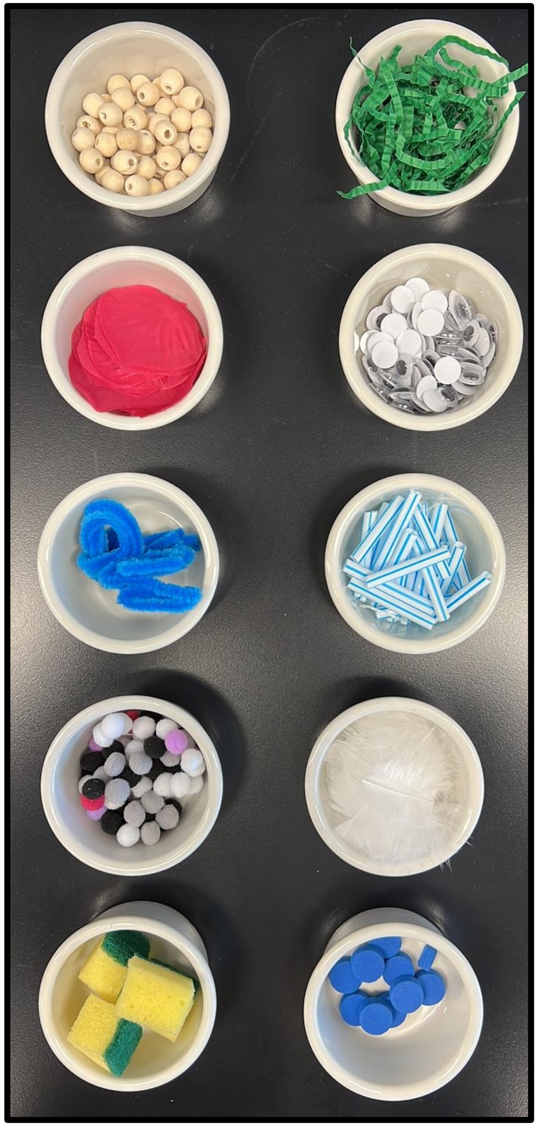

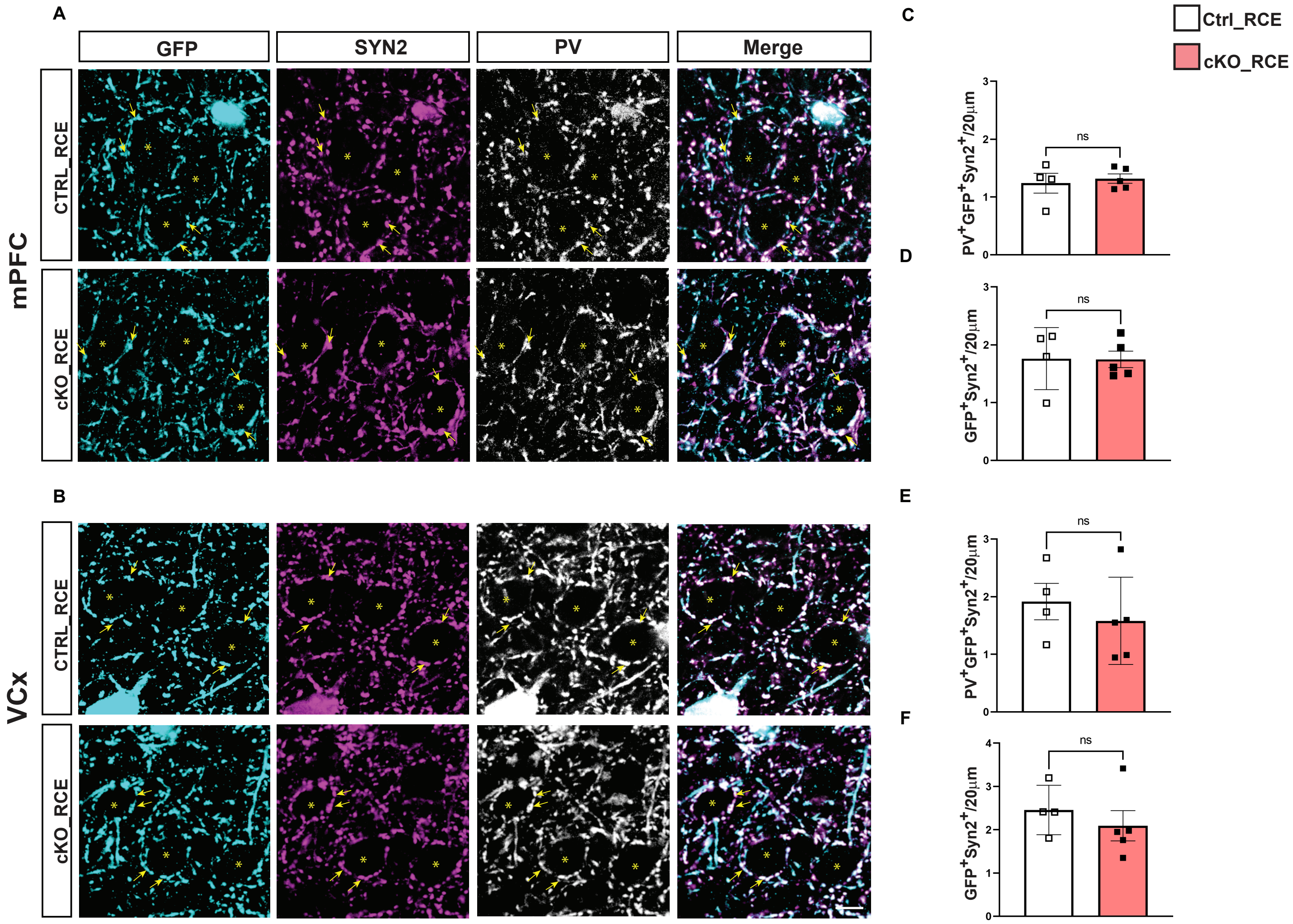
